# A factorial cluster-randomised controlled trial combining home-environmental and early child development interventions to improve child health and development: rationale, trial design and baseline findings

**DOI:** 10.1101/465856

**Authors:** Stella M Hartinger, Nestor Nuño, Jan Hattendorf, Hector Verastegui, Mariela Ortiz, Daniel Mäusezahl

## Abstract

**Background:** Exposure to unhealthy environments and poor cognitive development are the main risk factors that affect children’s health and wellbeing in low- and middle-income countries. Interventions that simultaneously address several risk factors at the household level have great potential to reduce these negative effects. We present the design and baseline findings of a cluster-randomised controlled trial to evaluate the impact of an integrated home-environmental intervention package and an early child development programme to improve diarrhoea, acute respiratory infections and childhood developmental outcomes in children under 36 months of age living in resource-limited rural Andean Peru.

**Methods:** We collected baseline data on children’s developmental performance, health status and demography as well as microbial contamination in drinking water. In a sub-sample of households, we measured indoor and personal 24-hour air concentration levels of carbon monoxide (CO) and fine particulate matter (PM_2.5_).

**Results:** We recruited and randomised 317 children from 40 community-clusters to four study arms. At baseline, all arms had similar health and demographic characteristics, and the developmental status of children was comparable between arms. The analysis revealed that more than 25% of mothers completed primary education, a large proportion of children were stunted and diarrhoea prevalence was above 18%. Fifty-two percent of drinking water samples tested positive for thermo-tolerant coliforms and the occurrence of *E.coli* was evenly distributed between arms. The mean levels of kitchen PM_2.5_ and CO concentrations were 213 μg/m^3^ and 4.8 ppm, respectively.

**Conclusions:** The trial arms are balanced with respect to most baseline characteristics, such as household air and water pollution, and child development. These results ensure the possible estimation of the trial effectiveness. This trial will yield valuable information for assessing synergic, rational and cost-effective benefits of the combination of home-based interventions.

**Trial registration:** retrospectively registered on 15^th^ January 2018. SRCTN reference: 26548981

## 1. Background

Children in low- and middle-income countries are frequently exposed to cumulative health and developmental risks often rooted in unhealthy environments [1]. Nearly 3 billion people worldwide still use solid fuels for cooking, a major source of household air pollution (HAP) [2]. HAP exposure drastically increment the risk of acute respiratory infections (ARI) [3], pneumonia [4] and other adverse outcomes such as cognitive deficits in children [5]. Eleven percent of the world population still rely on unimproved drinking water sources and 36% on unsafe sanitation [6]. Exposure to inadequate drinking water, sanitation and personal hygiene increases the risk for diarrhoea [7] and malnutrition [8]. Early child development (ECD) generates opportunities that shape child’s lifelong health and developmental status [9]. However, today two hundred million children worldwide are not developing their full cognitive potential [10]. This disparity has lifetime adverse consequences and impacts wellbeing of next generations [11].

Having access to safe drinking water (i.e. piped public taps) decreased school absenteeism [12]. Effective household water treatments combined with safe storage provided protection against diarrhoea [13]. Hand-washing with soap reduced diarrhoea by 40% [14]. The use of improved cookstoves (ICS) lessened HAP exposure and improved health [15]. ECD interventions enhanced health status [16] and development in children [16, 17].

The reduction of poor health and development requires integrated approaches addressing the underlying risks and structural determinants at the household level [18]. Multiple studies demonstrated that low-cost interventions focused on various health risks factors simultaneously had a synergic effect and increased opportunities for a sustained use over time [17, 19]. Integrating ECD and home-hygiene interventions is promising, as they both target young children within their homes [20]. The home-based approach enhances the effects of ECD interventions, which could also lead to improvements of children’s general health and affection, and mother’s mental health [21, 22]. Furthermore, home-based interventions can help reducing the use of material and human resources, providing a more holistic approach towards prevention.

In this article, we describe the design and baseline results of a cluster-randomised controlled trial seeking to develop an integrated cost-effective home-based package of environmental health and ECD intervention in rural Andean Peru.

## 2. Methods

### 2.1. Setting and subjects

The trial was conducted in the San Marcos and Cajabamba provinces, Cajamarca region, Andean Peru. We chose the locations based on the coverage of the national ECD programme (“Programa Nacional Cuna Mas” (PNCM)) and on-going relationships with local stakeholders who partook in a previous cluster-randomised controlled trial conducted in the region. The results of our previous research endeavours enhanced the implementation of this new trial (“IHIP-2 trial” in the following) [17, 20]. Both sites are high altitude rural resource-limited locations with chronic malnutrition and illiteracy [23]. The majority of the population are small-scale farmers living in 2-3 roomed houses with earthen floors and adobe walls, with traditional stoves or open fires for cooking [24].

### 2.2. Study design

We implemented a 2×2 factorial design trial applying two interventions individually and in combination: i) an environmental health package comprising a certified ICS, kitchen sink and hygiene education (IHIP); and ii) an early child development programme (ECD). This design led to four potential experimental conditions: i) IHIP & ECD (“IHIP+” in the following), ii) IHIP, iii) ECD and iv) Control. We chose this design because it randomises communities instead of individuals, avoiding contamination between participants.

We enrolled all families complying with the following inclusion criteria: i) had at least one child <1.5 months living at the household; ii) used solid fuels as main energy source for cooking/heating; iii) had access to piped water in the yard; iv) did not have moving plans for the next 24 months; and iv) did not participate in the PNCM.

### 2.3. Sample size

We calculated the sample size using the formula proposed by Hayes & Bennett for cluster-randomised trials [25], assuming three episodes of diarrhoea per child-year, a 25% reduction of the incidence compared to the control arm, and a coefficient of variation of 0.2. With 10 person-years of follow-up in each cluster, we calculated 16 clusters for the intervention and control arm to detect the anticipated reduction of incidence with a power of 80% at the 5% two-sided significance level. To account for potential loss to follow-up, we included 40 clusters with an average of 6.5 person-years of follow-up. For the ECD intervention, we used the ECD outcome (percentage of tasks solved above the mean of the study population) of our previous intervention study [17] and assumed 60% above mean for the intervention and 40% above mean in the control arm. Using the equivalent formula for proportions, we calculated that 15 clusters for intervention and control with ten children. Of note, the trial is sufficiently powered to compare each intervention against its control arm but does not allow pair wise comparisons among the four trial arms.

### 2.4. Recruitment

We carried out a census in 2015 to identify potential communities, children and pregnant women in their second and third trimester in collaboration with the Peruvian Ministry of Health (MINSA). Participants were enrolled between September 2015 and January 2016. If multiple eligible children were found in a household, we selected the youngest child.

### 2.5. Randomisation

We enrolled 82 eligible communities, which we aggregated into 40 community-clusters because of their partly close proximity to each other. After the enrolment, we allocated the communities into the four study arms using a covariate-based constrained randomisation as proposed by Moulton [26]. Based on geographic distance, we first divided the clusters into 8 strata of 4 clusters each and 1 stratum of 8 clusters. We then generated two million random allocation sequences and selected those for which the maximum difference between arms was i) ≤ 5 in terms of number of villages; ii) ≤10 for the number of children; iii) median village size (number of households) ≤10; iv) median altitude ≤250; v) proportion of households with a health post in the community ≤10 %-points; vi) proportion of households with a school within the community ≤10 %-points; and vii) proportion of households living in villages having an electricity connection ≤10 %-points. Of the 164 allocation sequences that fulfilled all criteria, one was randomly selected.

### 2.6. Design and implementation of study interventions

#### 2.6.1. The IHIP intervention

Between May and July 2015, we consulted communities on their preference on three preselected ICS. Forty-eight women were asked to cook with each ICS model for three consecutive days (9 in total). Participant women selected the ICS “OPTIMA” model and recommended further modifications before its installation. The modified stove was certified by the Peruvian national industrial certification authority (SENCICO)[27]. Kitchen sinks and stove parts were purchased locally to increase scalability. Both interventions were installed between November 2015 and February 2016.

#### 2.6.2. Early child development intervention

The ECD intervention was based on the PNCM [28]. The PNCM was launched in 2012 to improve the cognitive, social and emotional development of infants <3 years of age living in poverty. In addition, the programme sought to improve families’ knowledge and practices regarding caregiving, and strengthens the bond between mothers and children. The PNCM provides a home-visiting intervention in rural areas (“Acompañamiento a Familias” (AAF)) as opposed to a day-care service in urban settings. The AAF was implemented and evaluated in our trial.

Mother facilitators (MF) were trained women living in the participating communities and conducted weekly play-oriented, semi-structured activities with the participant mothers (or caretakers) and the child. MFs were selected using PNCM guidelines and received a one-day training session and monthly re-trainings. MFs were supervised by a technical assistant team (TA), which mentored and assisted MFs in the planning of the weekly sessions according to the PNCM guidelines. In addition, the TA team organised bi-monthly group sessions with participant mothers to share experiences, lessons or concerns of the ECD intervention. The TA team received a one-week training from PNCM experts in planning and delivering of the ECD sessions. All participating household received a set of age-specific toys to stimulate child’s psychomotor and cognitive development every two months (six packages in total). These educational materials aimed to foster communicative, socio-emotional and cognitive abilities and to teach mothers plays and games for daily interactions.

### 2.7. Primary and secondary outcomes

#### 2.7.1. Definition of the primary and secondary outcomes

We assessed childhood diarrhoea and ECD status as primary outcomes. We defined diarrhoea according to the World Health Organization (WHO) standards, passing of at least three loose stools within 24 hours [29], and ECD outcomes as an age standardised mean scores of psychomotor assessment (including socio-emotional, motor and cognitive skills, and communication abilities) in children <3 years of age.

The secondary outcomes included: i) ARI, defined as presence of cough and fever reported by the primary caretaker, according to the WHO standards [30]; ii) severe cases of diarrhoea defined as persistent diarrhoea for more than 14 days or bloody diarrhoea; iii) household and personal exposure to carbon monoxide (CO) and particle matter (PM_2.5_) in a sub-sample of 40 participants (“sentinel sub-sample” in the following); iv) presence of *E.coli* in drinking water samples; and v) compliance linked to the use of the interventions. We assessed compliance on stove and sink use and hygiene through spot check observations, 24-hour recall data and direct observations in the sentinel sub-sample. ECD compliance was defined as the total number of MFs visits and mother’s satisfaction associated to the MFs visits.

#### 2.7.2. Data collection of primary and secondary outcomes

We carried out active and passive surveillance to collect diarrhoea and ARI morbidity data during the 12-month follow-up. The fieldworker team (FW) visited all households’ weekly and collected daily and weekly information from the mother or caretaker about the occurrence of signs and symptoms of child diarrhoea and ARI. The FW was instructed to obtain two measurements of respiratory rates, which increased the specificity for a diagnosis without a loss of sensitivity [31]. To define the severity of the disease, for diarrhoea we collected additional information on observed blood in the stools. For ARI, we measured respiratory rate, heart rate and oxygen saturation in blood (SpO_2_) with portable pulse oximeters (PPOs) (Masimo iSpO_2_ Rx) and multisite reusable sensors (Masimo M-LNCS YI SpO_2_) connected to tablets (Lenovo TAB 2 A7-10). We used the *lambdanative* framework to provide a real-time diagnostics of ARI and its severity [32]. Respiratory rate was recorded using the RRate app module [33]. Severely ill children were referred to the local healthcare facility for further evaluation. The FW received a five-day initial training for morbidity data collection with monthly re-training sessions of two hours. All the surveillance devices and tools were tested on a daily-basis between February and April 2016. In addition, the passive surveillance team (PS) collected health data from our study participants monthly at their local community-based health centres (22 in total). To ensure that they were also collecting SpO_2_ in each evaluation, we provided PPOs, sensors and tablets to local health centres. During the visits, the PS also trained local health personnel on the maintenance and use of the PPOs. Anthropometric measurements were collected from participant’s clinical records. We assessed stunting and underweight following the WHO standards [34].

The environmental team (EV) collected household air pollution and drinking water samples. The EV received a seven-day initial training and monthly re-training sessions. We collected 24-hour kitchen and personal CO and PM_2.5_ exposure data. We installed one EL-USB-CO (LACAR Electronics) monitor and one HAP measuring device for indoor use APROVECHO-5000 at a one-meter distance from the ICS and at standard breathing height (1.5 meters). To measure personal exposure, mothers were carrying a vest equipped with one EL-USB-CO (LASCAR Electronics) monitor and one micro personal aerosol exposure monitor (RTI-INTERNATIONAL) fitted. We asked mothers to wear the vest for 24 hours only taking them off for sleeping and personal hygiene.

We obtained drinking water samples from the child’s main drinking source, and then transported to the field station’s laboratory in San Marcos in cooled thermal bags. At the laboratory, the samples were analysed for thermotolerant (faecal) coliforms using a membrane-filtration method from the Oxfam DelAgua water testing kit [35]. All yellow colonies forming units were considered positive for *E.coli* growth. These were then collected and placed in transportation vials media and send to Lima for further phenotypic and genotypic identification and antibiotic resistance testing.

HAP data were obtained from the sentinel sub-sample five times (before and after ICS installation and three times during follow-up). Water samples were collected at the baseline and end of study for all study participants and in the sentinel sub-sample four additional times over the follow-up.

To assess ECD status, the TA carried out an assessment using the nationally validated Peruvian Infant Development Scale (ESDI) tool [36] at baseline and end of study. The TA received a one-week training from PNCM experts. For evaluating ECD improvements, the TA also implemented the ESDI tool at end of study. In parallel, a group of specialised psychologists conducted the Bayley Scales of Infant and Toddler Development (BSID) [37] instrument for comparability.

Finally, the capacity building team (CBT) was responsible for re-training participants on hygiene, hand-washing, boiling practices and ICS maintenance. The CBT conducted monthly reinforcement visits and collected data on the IHIP intervention condition. A local ICS constructor was hired in case additional maintenance of the intervention was required. During their visits, the FW, CBT and TA conducted regular spot check observations and collected maternal reports on the usage and quality of the interventions as well as household and environment hygiene.

#### 2.7.3. Socio-economic survey

The FW implemented a socio-economic questionnaire at baseline and end of study. The objective was to assess household demographics, education and economic characteristics, general stove use, and household water management. We conducted the baseline assessment between September 2015 and February 2016. The FW received a one-week of training.

#### 2.7.4. Data quality management

The field coordinator team revised information collected on a daily basis to reduce the chance of missing data. They also trained the field staff, double-checked questionnaires and conducted regular household quality visits monthly. Ten percent of all data collected in the trial was double entered to ensure data quality. To reduce the possibility of courtesy bias, household data collection routes of FW were changed every two months.

## 3. Results

### 3.1. Enrolment

From the screening census, we identified 102 communities with 574 potential children. During enrolment we found that 237 families were no longer eligible because i) they participated in another social programme (N=167); ii) they did not fulfil the inclusion criteria (N=34), and iii) they rejected participation (N=36). We re-enrolled between January and February 2016 because 28 children were not available or rejected to participate in the project at the beginning of the follow-up. In total, 317 households in 10 clusters per arm participated in the trial (Figure 1).

**Figure 1:**
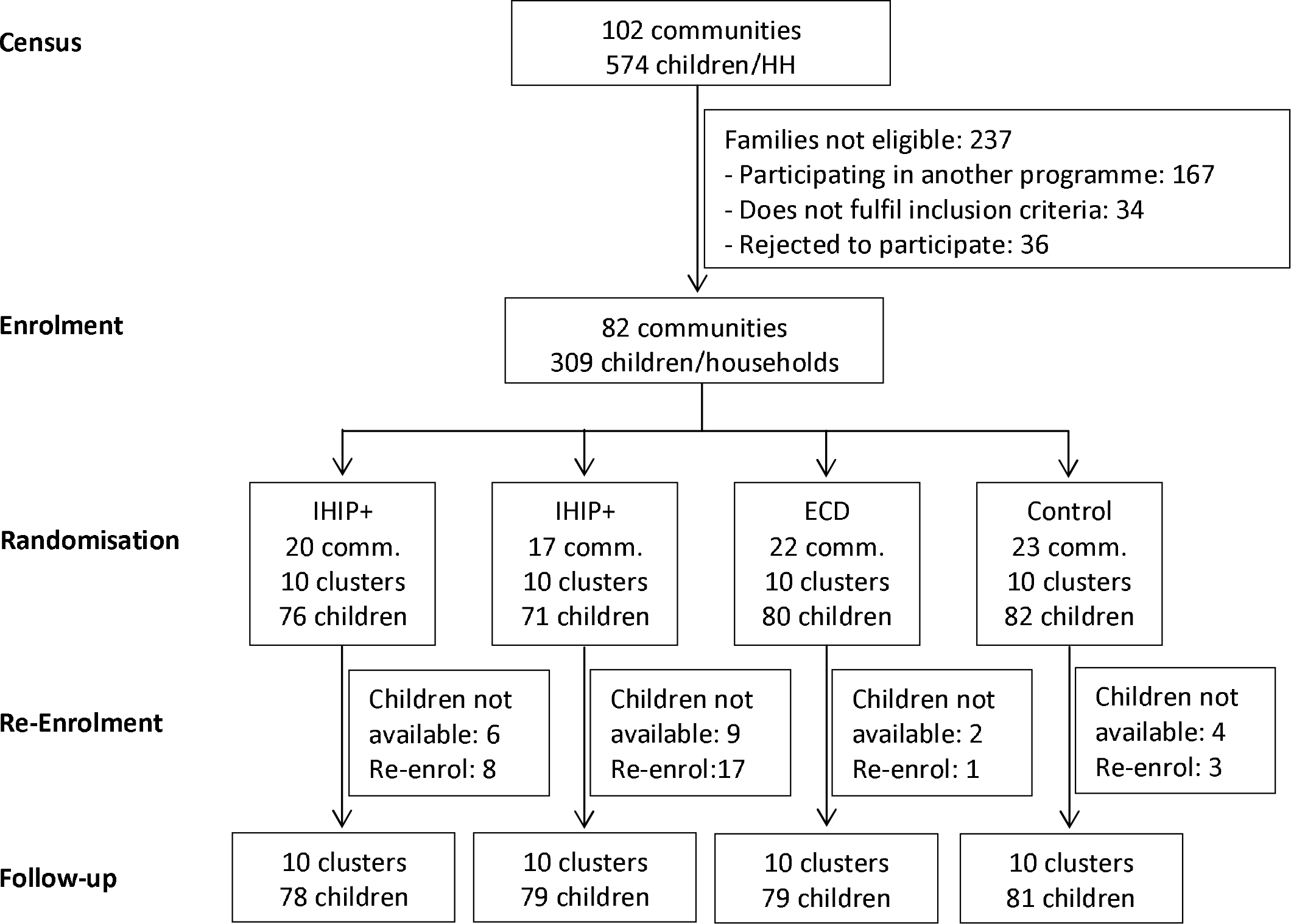
Flow of participants.

### 3.2. Baseline characteristics

Baseline analysis included the demographic and health characteristics, the household drinking water quality and levels of ECD indicators for each participant. We measured household and personal air pollution concentrations in the sentinel sub-sample.

#### 3.2.1. Demographic and household characteristics

The four arms were balanced in most of the domains (Table 1). Mothers from the four arms were of similar mean age, had similar levels of primary education and years of schooling. The majority of household had adobe walls, earthen floors, tiled roofs, and piped water system. We observed differences among trial arms in the proportion of children below 1 year. This imbalance disappears comparing *IHIP versus* no-IHIP or ECD *versus* no-ECD arms. We also observed imbalance in the proportion of households with electricity. Although living in villages having an electricity connection was a randomisation constraint it did not ensure balance at household level (in villages with connection the proportion of households with electricity varied from 50 to 100%).

**Table 1.**
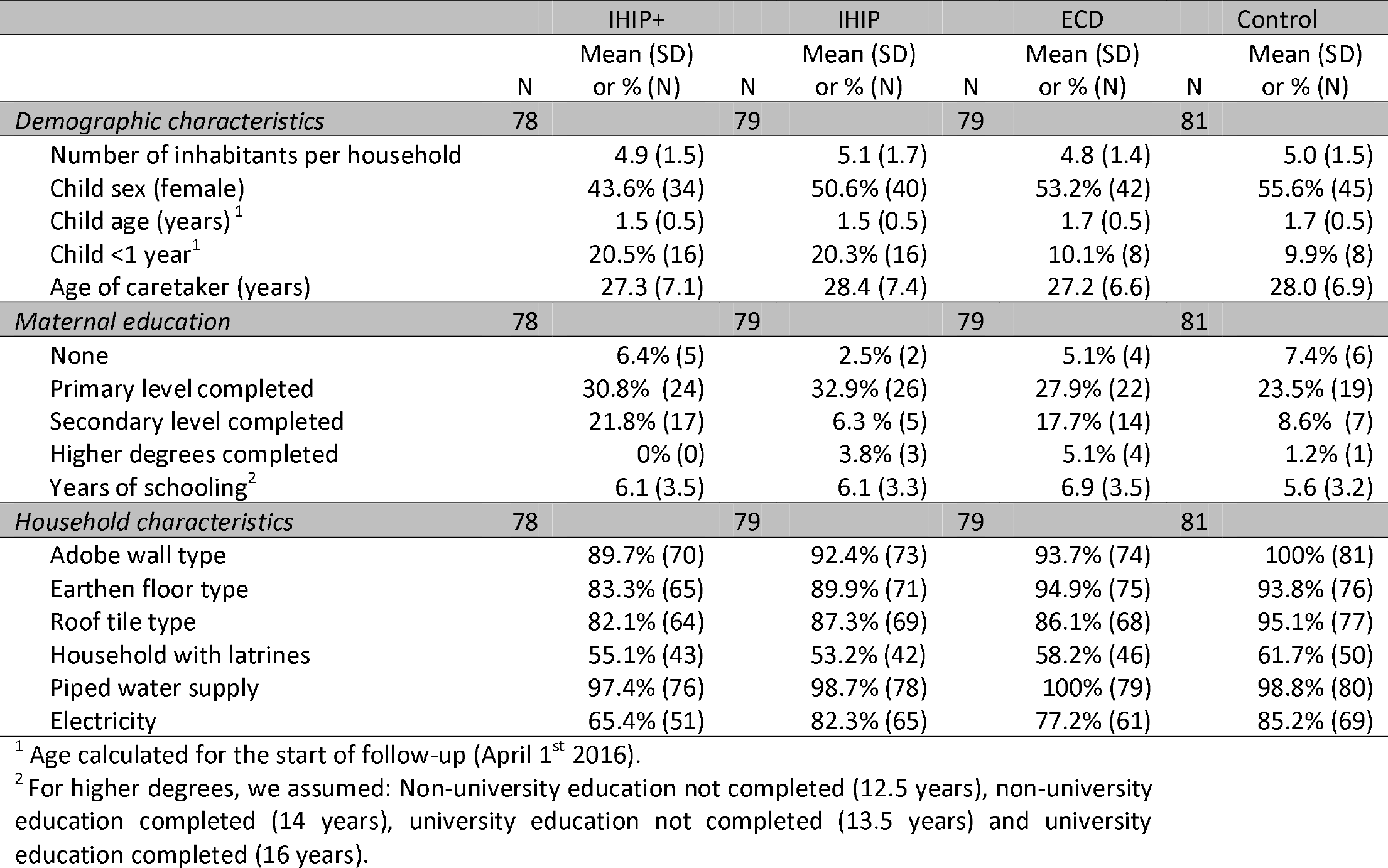
Demographic and household characteristics of households in rural Andean Peru.

#### 3.2.2. Health characteristics

Children in all arms had similar weight at birth. Imbalance was observed in the prevalence of stunting, with the highest proportion in the ECD arm (36.9%). Two-week diarrhoea prevalence was balanced between arms, as were two-week prevalence of cough and fever. However, we did observed differences between the health insurance coverage between arms, with the highest lack of coverage in the IHIP+ (10.3%) and the lowest in the control arm (1.3%). Vaccination coverage, varied between arms (Table 2). The highest complete doses of vaccines for children between 6-11 months were in the ECD, and the lowest in the control arm (40%). These percentages change when we look at children between 12-24 months, where coverage levels in the ECD arm increase up to 70% and the four arms are balanced. SpO_2_ and respiratory rate levels were similar in all arms (Table 2).

**Table 2.** Children’s health status and vaccination coverage in rural Andean Peru.

|  | IHIP+ |  | IHIP |  | ECD |  | Control |  |
| --- | --- | --- | --- | --- | --- | --- | --- | --- |
|  | Mean (SD) |  | Mean (SD) |  | Mean (SD) |  | Mean (SD) |  |
|  | N | or % (N) | N | or % (N) | N | or % (N) | N | or % (N) |
| **Anthropometrics<sup>1</sup>** | 78 |  | 79 |  | 79 |  | 81 |  |
| Height-for-age, Z-values | 64 | -1.3 (1.1) | 50 | -1.0 (0.9) | 64 | -1.5 (1.1) | 61 | -1.4 (0.8) |
| Stunting | 64 | 17.2% (11) | 50 | 9.8% (5) | 64 | 36.9% (24) | 61 | 18.0% (11) |
| Weight-for-age, Z-scores | 64 | -0.4 (1.0) | 51 | -0.1 (0.8) | 65 | -0.6 (0.9) | 61 | -0.5 (0.9) |
| Underweight | 64 | 6.3% (4) | 51 | 2.0% (1) | 65 | 10.8% (7) | 61 | 4.9% (3) |
| **Children's health status** | 78 |  | 79 |  | 79 |  | 81 |  |
| Weight at birth |  | 3.1 (0.6) |  | 3 (0.5) |  | 3.1 (0.5) |  | 3.1 (0.4) |
| Diarrhoea, two-weeks prevalence |  | 18.4% (14) |  | 22.8% (18) |  | 19% (15) |  | 26.3% (21) |
| Fever, two-weeks prevalence |  | 21.8% (17) |  | 21.5% (17) |  | 25.3% (20) |  | 23.5% (19) |
| Cough, two-weeks prevalence |  | 35.1% (27) |  | 31.7% (25) |  | 39.2% (31) |  | 29.1% (23) |
| Children with no health coverage |  | 10.3% (8) |  | 10.1% (8) |  | 5.1% (4) |  | 1.2% (1) |
| **Children's vaccination** | 78 |  | 79 |  | 79 |  | 81 |  |
| Tuberculosis (BCG) | 77 | 89.6% (69) |  | 91.1% (72) |  | 94.9% (75) |  | 95.1% (77) |
| Measles | 77 | 50.7% (39) | 78 | 60.3% (47) | 77 | 67.5% (52) |  | 64.2% (52) |
| Diphtheria, convulsive cough and tetanus (DTP)* |  |  |  |  |  |  |  |  |
| 1<sup>st</sup> dose | 77 | 96.1% (74) |  | 94.9% (75) |  | 98.7% (78) |  | 95.1% (77) |
| 2<sup>nd</sup> dose | 73 | 90.4% (66) | 77 | 89.6% (69) |  | 97.5% (77) | 79 | 93.7% (74) |
| 3<sup>rd</sup> dose | 64 | 87.5% (56) | 69 | 79.7% (55) | 72 | 93.1% (67) | 73 | 82.2% (60) |
| Polio (OPV)** |  |  |  |  |  |  |  |  |
| 1<sup>st</sup> dose | 77 | 96.1% (74) |  | 96.2% (76) |  | 98.7% (78) |  | 96.3% (78) |
| 2<sup>nd</sup> dose | 73 | 89.0% (65) | 77 | 90.9% (70) |  | 97.5% (77) | 79 | 93.7% (74) |
| 3<sup>rd</sup> dose | 64 | 78.1% (50) | 69 | 79.7% (55) | 72 | 90.3% (65) | 73 | 75.3% (55) |
| Influenza | 77 | 62.3% (48) |  | 63.3% (50) |  | 81.0% (64) |  | 69.1% (56) |
| Hepatitis B | 77 | 62.3% (48) |  | 54.4% (43) |  | 79.8% (63) |  | 69.1% (56) |
| **Recommended vaccines 6-11 months** | 23 |  | 28 |  | 24 |  | 15 |  |
| All vaccines<sup>2</sup> |  | 69.6% (16) |  | 60.7% (17) |  | 87.5% (21) |  | 40% (6) |
| No vaccines |  | 0% (0) |  | 7.1% (2) |  | 0% (0) |  | 0% (0) |
| **Recommended vaccines 12-24 months** | 41 |  | 41 |  | 48 |  | 58 |  |
| All vaccines<sup>3</sup> |  | 68.3% (28) |  | 70.7% (29) | 46 | 84.8% (39) |  | 70.7% (41) |
| No vaccines |  | 2.4% (1) |  | 4.9% (2) |  | 2.1% (1) |  | 0% (0) |
| **Portable pulse oximetry assessment<sup>4</sup>** | 64 |  | 59 |  | 63 |  | 68 |  |
| Oxygen saturation (SpO<sub>2</sub>) |  | 93.5 (2.8) |  | 94.2 (2.5) |  | 93.8 (3.8) |  | 93.8 (2.8) |
| Respiratory rate per/min. |  | 27.7 (3.7) |  | 26.4 (3.0) |  | 27.2 (2.7) |  | 27.3 (3.1) |

<sup>1</sup>As estimated on 15<sup>th</sup> December 2015 (±31 days). If several estimates within the range were available, the one closest to 15<sup>th</sup> of December was selected.
\*DTP vaccination: 1<sup>st</sup> dose: 2 month; 2<sup>nd</sup> dose: 4 month; and 3<sup>rd</sup> dose: 6 month.
\*\* Polio vaccination: 1<sup>st</sup> dose: 2 month; 2<sup>nd</sup> dose: 4 month; and 3<sup>rd</sup> dose: 6 month.
<sup>2</sup> It includes BCG (one dose), DTP (three doses) and OPV (three doses).
<sup>3</sup> It includes BCG (one dose), DTP (three doses), OPV (three doses) and measles (one dose).
<sup>4</sup> Results from children with no observable danger signs, fever, cough and difficulty to breath at the time of the assessment.

### 3.3. Household microbial contamination

We obtained 314 water samples. Some 52.9% (N=166) tested positive for thermo-tolerant bacteria. The number of positives and the occurrence of *E.coli* and Enterobacter were evenly distributed between arms. We observed imbalances in the occurrence of *Klebsiella* and *Citrobacter*, although the number of cases are small (Table 3).

**Table 3.**
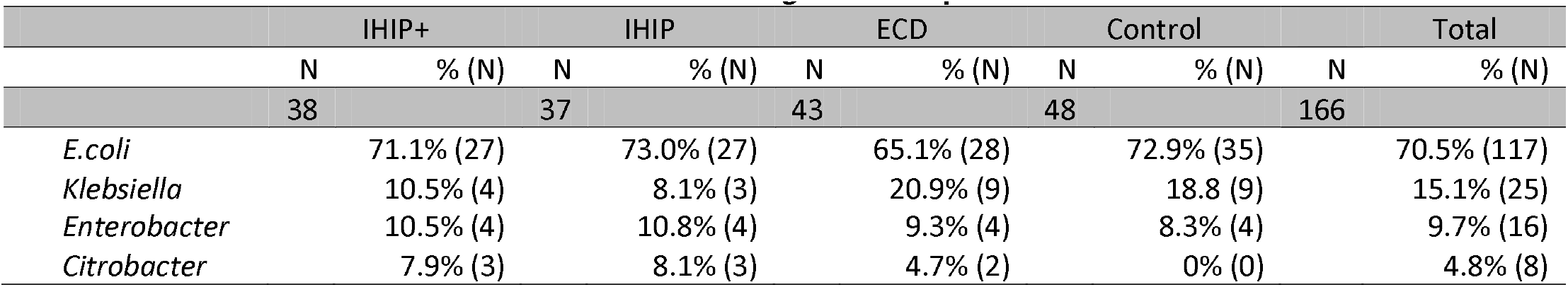
Thermo-tolerant bacteria from household drinking water samples in rural Andean Peru.

### 3.4. Household and personal air pollution

We measured indoor PM_2.5_ and CO concentrations stationary from the kitchen environments and personal exposure (CO only) before the installation of the ICS. Data was collected from 33 households. The average household PM_2.5_ and CO concentration were 213 µg/m^3^ (SD=166.1) and 4.8 ppm (SD=3.7) respectively (Table 4).

**Table 4.** HAP measurements at baseline in rural Andean Peru.

|  | N | Mean (95% IC) | Median (95% IC) | Geometric mean (95% IC) |
| --- | --- | --- | --- | --- |
| *Kitchen* | 33 |  |  |  |
| PM<sub>2.5</sub> (µg/m<sup>3</sup>) |  | 213 (154.9 – 271.2) | 46.5 (23.5 – 64.6) | 48.1 (37.5 – 61.7) |
| CO (ppm) |  | 4.8 (3.5 – 6.2) | 1.1 (0.7 – 1.7) | 1.3 (1.0 – 1.7) |
| *Personal* | 33 |  |  |  |
| CO (ppm) |  | 1.4 (0.8 – 2.0) | 0 (0 – 0) | 2.4 (1.9 – 3.1) |

### 3.5. Early child development assessment

Child psychomotor and cognitive development indicators of 305 study children indicated similar performance in all developmental domains across arms (Figure 2).

**Figure 2.**
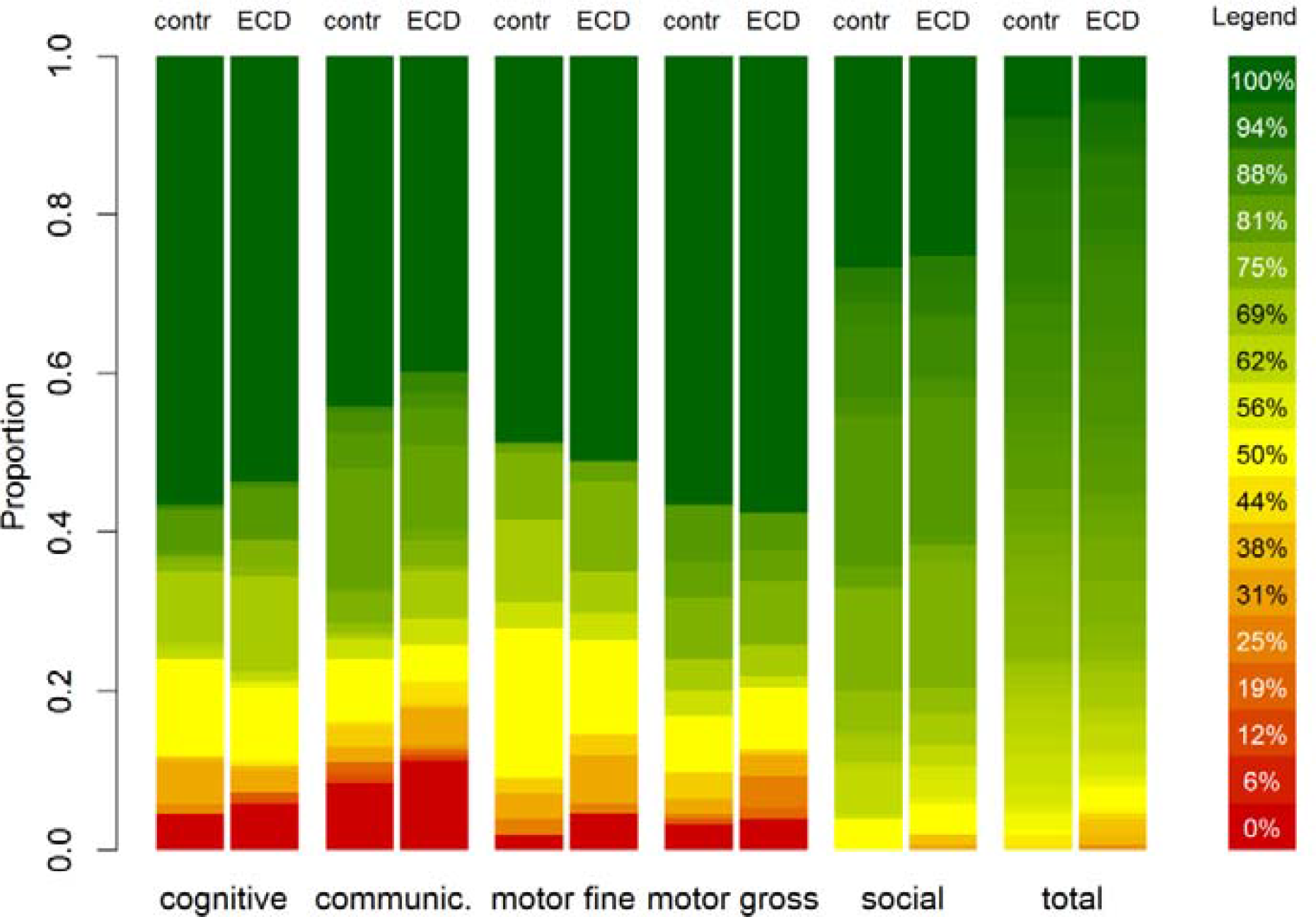
Proportion of tasks completed for all ECD-developmental domains.

## 4. Discussion

We present the baseline results of a cluster-randomised trial evaluating an integrated home-environmental intervention package and an ECD programme to improve diarrhoea, ARI and childhood developmental outcomes in children under 36 months of age living in resource-limited rural Andean Peru. This trial uses a robust 2×2 factorial design that allows the assessment of two interventions in a single study without increasing sample size [38]. For the ECD intervention, we implemented one component of the Peru’s national ECD programme (PNCM).

The trial included 317 rural families living in 82 communities from two Peruvian provinces. Baseline results indicate the trial arms are balanced with respect to most baseline characteristics, but given the limited number of clusters and the amount of characteristics presented, we also found a few imbalances. Imbalances observed will be considered in the primary trial analysis comparing each intervention separately with its counterfactual, i.e. IHIP versus no-IHIP to assess the impact of the IHIP intervention, and ECD *versus* no-ECD arms to evaluate the effects of the ECD intervention. While the ECD performance measurements at baseline appear to be inaccurately high, results are robust to no difference in ECD status between trial arms. We also found that households across all arms share a notable burden of household environmental risks. Microbial contamination of drinking water does not comply with the Peruvian and WHO standards of zero viable coliforms in potable water samples [39, 40]. Indoor and personal 24-hour CO air pollution measurements in our study meet WHO guidelines [41], but PM2.5 levels exceed the threshold of 25 µg/m^3^ recommended by the WHO on average 8 times (213 µg/m^3^) [42].

There is an important need for implementing effective national ECD programmes to address the burden of poor development in early childhood [43]. Experiences demonstrate that implementing ECD programmes in low- and middle-income countries is feasible and cost-effective [43, 44]. Our trial offers an ideal opportunity to generate strong evidence regarding the PNCM-programme’s efficacy and support the Peruvian national government in its expansion on a wider scale.

The use of active surveillance in this trial will maximize sensitivity while passive surveillance increases the likelihood that true cases will be diagnosed, increasing specificity [45]. To synchronise ARI diagnosis with new national guidelines (stipulated by the MINSA) and address potential limitations of passive ARI surveillance, we equipped all the health centres in our working area with PPOs. Both, health and project field staff were trained, monitored and supervised monthly in the use of PPOs to assess ARI cases equally. PPOs have been underutilised in resource-limited health care settings for a variety of reasons, including high cost, inadequate supply and lack of training [46]. Our design overcomes these limitations and enhances the research of the usefulness and effectiveness of PPOs for ARI case assessments. In addition, PPOs show to be optimal tools for identifying children with hypoxemia, an indicator for severe respiratory diseases [47]. Previous research indicated that the reference SpO_2_ thresholds for hypoxemia are lower at high altitudes in comparison to those at sea level [48]. A median SpO_2_ hypoxemia threshold of 96% for children below 5 years living at 2500 m.a.s.l has been proposed [49]. Our SpO_2_ results in healthy children living between 2250-3900 m.a.s.l are lower than the suggested threshold. Hence, pondering specific SpO_2_ hypoxemia thresholds for children living in high altitudes becomes a challenge. Our systematic health data collection using PPOs may further pave the way for assessing SpO_2_ hypoxemia threshold values at high altitude settings for children with and without related ARI symptoms [50].

Little attention has been paid to user’s perspectives and preferences in the design of ICS interventions in the past [51]. We conducted a community consultation to select an ICS model that was both locally accepted and efficient. According to recent evidence, those who participate in community consultations to select ICS models have a higher odd of increasing ICS use over time [52].

Our study has some limitations. Because of the nature of the trial design, interventions could not be blinded. The use of tablet-based technologies to measure pulse oximetry was a challenge for both, health centre and field staff. We trained the health centre personnel on the correct use and benefits of introducing PPOs in their daily work in two group sessions. However, due to frequent changes of health centre’s personnel, we had to retrain individually new staff on-site on a monthly basis. Baseline ECD evaluation and the age-specific assessment tools were not as straight forward to apply as we had experienced previously [17]. Despite the assertion from the PNCM programme experts that also novices could apply the ESDI tool, we found that the lack of familiarity and experience among the field staff consistently produced unusually high scores in all individuals (at baseline).

However, in the context of this baseline assessment, the screening was conduct to assess the relative ECD status between study arms as opposed to estimate absolute differences in ECD psychomotor and cognitive indicators. Hence, overall high scores do not invalidate our conclusion that randomisation was successful given study arms had similar ECD levels at baseline. To diminish any further potential bias at the end of study, we will apply both the ESDI and BSID tools.

We believe despite the limitations, the IHIP-2 trial will generate needed evidence on the potentially synergistic benefits of combining ECD and environmental health interventions. We actively sought to involve national and governmental actors (i.e. PNCM, SENCICO, MINSA) in developing our interventions and throughout the study to foster sustainability and reinforce collaborative engagements in the future.

## 5. Conclusion

In this paper, we present the baseline results of a factorial cluster-randomised trial evaluating an integrated home-environmental intervention package and an early child development intervention in children under 36 months of age living in rural Andean Peru. The trial arms were balanced with respect to most baseline characteristics, air and water contamination and child’s developmental status. Baseline results determine that the trial’s randomisation was successful and study arms are comparable for analysis at the end of study. The results of this trial will yield valuable information for assessing synergic, rational and cost-effective benefits of the combination of home-based interventions.

## 6. List of abbreviations

AAF: Acompañamiento a Familias
ARI: Acute respiratory infections
BSID: Bayley Scales of Infant and Toddler Development tool
CBT: Capacity building team
CO: Carbon monoxide
ECD: Early child development
ESDI: Peruvian infant development scale tool
EV: Environmental team
FW: Fieldworker team
HAP: Household air pollution
ICS: Improved cookstoves
IHIP: Integrated home-based intervention package
MINSA: Peruvian Ministry of Health
MF: Mother facilitators
PM_2.5_: fine particulate matter
PNCM: Program Nacional Cuna Mas
PPO: Portable pulse oximeters
PS: Passive surveillance team
SENCICO: Peruvian national industrial certification authority
SpO_2_: Oxygen saturation in blood
TA: Technical assistant team
WHO: World Health Organization

## 7. Declarations

### 7.1. Ethics approval and consent to participate

The study (trial registry: ISRCTN-26548981) was approved by the Cajamarca Regional Health Authority and the Universidad Peruana Cayetano Heredia (UPCH) (Ref 268-12-15). Community leaders and local authorities from the study area signed a collaborative agreement with the UPCH before study implementation. Families who agreed to participate in the study signed an informed consent. No incentives were given to foster participation.

### 7.2. Consent for publication

Not applicable

### 7.3. Availability of data and material

The datasets used and/or analysed during the current study are available from the corresponding author on reasonable request.

### 7.6. Authors’ contributions

DM and SH designed the study and obtained the funding; SH, NNM and HV collected data; SH, NNM, HV and MO were in charge of implementing interventions; JH provided with essential statistical advice and with HV generated the randomisation for the study. SH, HV, NNM, JH analysed and interpreted the data; SH wrote the first draft manuscript; DM, HV, NNM, JH and MO interpreted the data, performed critical revisions of the manuscript and contributed to the writing; SH, DM, HV, MO provided administrative, technical and material support; SH, NNM and DM, coordinated and supervised the study.

## 7.7. Acknowledgements

The authors would like to express their appreciation to the study families for their kind participation and the local authorities for their continuous support. We also express our gratitude to the field coordinators, especially to Mrs. Angelica Fernandez and Ms. Maria Luisa Huaylinos, for their unfailing support.

